# The spontaneous neoantigen-specific CD4^+^ T cell response to a growing tumor is functionally and phenotypically diverse

**DOI:** 10.1101/2025.03.25.645281

**Authors:** Ryan Q Griswold, Spencer E Brightman, Karla Soria Zavala, Manuel Azaid Ordaz-Arias, Navid Djassemi, Rukman R Thota, Martin S Naradikian, Hannah Dose, Suzie Alarcon, Vijayanand Pandurangan, Bjoern Peters, Aaron M Miller, Ezra E W Cohen, Stephen P Schoenberger

**Affiliations:** Center for Cancer Immunotherapy, La Jolla Institute for Immunology, La Jolla, California, USA; Biomedical Sciences Program, School of Medicine, University of California San Diego, La Jolla, CA, USA; Division of Vaccine Discovery, La Jolla Institute for Immunology, La Jolla, California, USA; AUGenomics, San Diego, CA; La Jolla Institute for Immunology, La Jolla, California, USA; Division of Hematology and Oncology, University of California San Diego Moores Cancer Center, La Jolla, California, USA; Department of Medicine, University of California San Diego, La Jolla, California, USA

## Abstract

CD4^+^ T cells play critical roles in the positive and negative regulation of cellular immunity through the many functional subsets they comprise. The progressive growth of immunogenic tumors which nonetheless generate mutation-specific T cells suggests that effective immune control may be avoided or suppressed at the level of the neoantigen-specific CD4^+^ T cell response. We used a tetramer specific for a validated neoantigen, CTLC_H129>Q_/I-E^k^, to characterize the ontogeny of natural CD4^+^ T cell responses to an aggressive and poorly immunogenic Major Histocompatibility Complex Class II (MHCII)-deficient tumor, SCC VII, during progressive growth or following therapeutic peptide vaccination. We find that the natural CD4^+^ T cell response to a growing tumor is phenotypically and functionally diverse, with distinct subsets including type 1 helper (T_h_1), T follicular helper (T_fh_)-like, and regulatory T cell (T_reg_) lineages appearing as early as 9 days after tumor implantation. Therapeutic vaccination using the CLTC_H129>Q_ peptide in adjuvant plus α-PD-1 sharply reduces the frequency of CLTC_H129>Q_-specific T_reg_ frequency in both tumor and tumor-draining lymph node (tdLN). Single cell transcriptomic analysis of CLTC-specific CD4^+^ T cells recapitulated and extended the diversity of the response, with TCRs of varying affinity found within each functional subset. The TCR affinity differences did not strictly correlate with function, however, as even the lowest affinity TCRs isolated from T_reg_ can mediate therapeutic efficacy against established tumors in the setting of adoptive cellular therapy (ACT). These findings offer unprecedented insight into the functional diversity of a natural neoantigen-specific CD4^+^ T cell response and show how immunotherapeutic intervention influences the phenotype, magnitude, and efficacy of the anti-tumor immune response.

****What is already known on this topic**:** Little is known about the ontogeny, architecture, development of the CD4^+^ NeoAg-specific repertoire induced by progressively-growing tumor. This study was performed to address this topic and contribute new information to aid in its understanding

****What this study adds**:** This study reveals that the NeoAg-specific CD4^+^ T cell response to a growing tumor is phenotypically and functionally diverse, featuring a range of functional T cells subsets including T_H_1, T_FH_, and T_reg_ expressing a range of functional TCR avidities, and demonstrates how an immunotherapeutic NeoAg vaccine can alter their relative composition within the tumor and tumor-draining lymph node.

****How this study might affect research, practice or policy**:** This study offers new insights into the diversity of NeoAg-specific CD4^+^ T cells and their response to a tumor in the presence or absence of immunotherapeutic intervention. This information could lead to new approaches to immune monitoring in the clinical setting of checkpoint blockade immunotherapy and cancer vaccines. Furthermore, we show that T_reg_ can be a potent source of TCRs that can mediate therapeutic benefit in the setting of adoptive cell therapy (ACT).

## Background

Natural NeoAg-specific T cells are associated with durable remission in some cancers and are crucial to the success of immunotherapy in many others, including those mediated by immune checkpoint blockade, personalized cancer vaccines, and some adoptive cell therapies (1–7). Whereas the contribution of NeoAg-specific CD8^+^ T cells in controlling MHC-I-expressing tumors is well documented in a growing body of literature, the role of NeoAg-specific CD4^+^ T cells is less well understood (8–11). This is likely due in part to the restricted expression of MHC-II which is normally limited to professional antigen-presenting cells (pAPC) such as dendritic cells (DC), B cells, and macrophages (MΦ), but can be either canonically expressed in some tumor types (e.g. B-cell lymphomas) or induced in others (e.g. melanomas) (12). Recent evidence suggests that MHC-II expression does not necessarily enable a tumor to directly present endogenous NeoAgs that can nonetheless access the class II presentation pathway or that MHCII^+^ tumor cells may not act as de facto APCs, potentially due to a lack of costimulatory molecules like CD80 (13, 14).

Depending on the cytokine, costimulatory, and TCR signals present during their initial priming, CD4^+^ T cells can develop into a range of specialized functional subsets (15–17). These include type 1 helper (Th1) cells which secrete pro-inflammatory cytokines and promote cellular immunity, type 2 helper (Th2) cells which secrete anti-inflammatory cytokines and promote humoral immunity, Tfh cells that aid the development and function of B cells, and inhibitory subtypes like Tregs and Type 1 regulatory (Tr1) cells (18–20).

NeoAg presentation to CD4^+^ T cells by pAPCs can produce a variety of therapeutic benefits in cancer. These include pAPC activation via CD40:CD40L which can result in enhanced priming of effector and memory CD8^+^ T cells by DCs, acquisition of a tumoricidal phenotype in MΦ, and induction of class switching and affinity maturation in B cells (21–25). NeoAg-specific CD4^+^ T cells can also secrete inflammatory cytokines such as IFN-γ and TNF within the tumor microenvironment (TME), which can directly inhibit tumor growth, suppress pro-tumor accessory cells, and inhibit tumor vasculature (26).

Given the range of potent effector pathways available to NeoAg-specific CD4^+^ T cells for the control of incipient cancers, the progressive outgrowth of primary and metastatic tumors suggests that these either do not develop or are actively suppressed under physiologic conditions. The former could involve a failure at any number of levels in the generation of CD4^+^ T cells including antigen presentation, engagement of the proper TCR repertoire, clonal expansion, or effector development. The latter could involve various immunosuppressive mechanisms, including the action of tumor specific Tregs or deviation into functional subsets that are unable to mediate tumor control. Distinguishing among these possibilities has been difficult as the developmental course of NeoAg-specific CD4^+^ T cells responding to progressive tumor growth *in vivo* is largely unknown in terms of phenotype, magnitude, and TCR repertoire.

We have addressed these questions in an aggressive low tumor mutational burden (TMB) preclinical tumor model SCC VII which generates CD4^+^ and CD8^+^ T cells specific for a set of 5 NeoAg following whole cell vaccination which can be engaged by a therapeutic peptide vaccine which requires both sets for its efficacy (27). Using CLTC_H129>Q_/I-A^k^ tetramers to isolate and characterize Cltc_H129>Q_-specific CD4^+^ T cells during progressive growth of SCC VII tumors, our data reveal that the NeoAg-specific CD4^+^ T cell response is functionally and phenotypically diverse, containing Treg, Tfh-like, and Th1 cells expressing TCRs of varying affinity within its repertoire. The response is also dynamic, as the relative ratios of these subsets changes over time and in response to vaccination. This provides an unprecedented insight into the ontogeny of the natural spontaneous NeoAg-specific CD4^+^ T cells responding to progressive tumor growth in the absence of other manipulation and how therapeutic NeoAg vaccination influences this development.

## Results

### Therapeutic Vaccination Increases the Magnitude of the NeoAg-specific CD4^+^ T cell Response

The parental SCC VII line grows aggressively following subcutaneous (s.c.) inoculation in untreated C3H mice (Figure 1a). Treatment with either α-PD-1 or a NeoAg vaccine consisting of a 20mer peptide containing the CLTC_H129>Q_ (Mut_48) epitope plus poly(I:C) beginning 6 days post tumor inoculation did not substantively impede tumor growth, whereas combining these modalities significantly reduced tumor development. The efficacy of this combined immunotherapy partially depends on CD4^+^ T cells and their ability to engage CD40, as either CD4^+^ T cell depletion or CD40L blockade after initiating treatment hindered its efficacy (Figure 1b).

**Fig 1:**
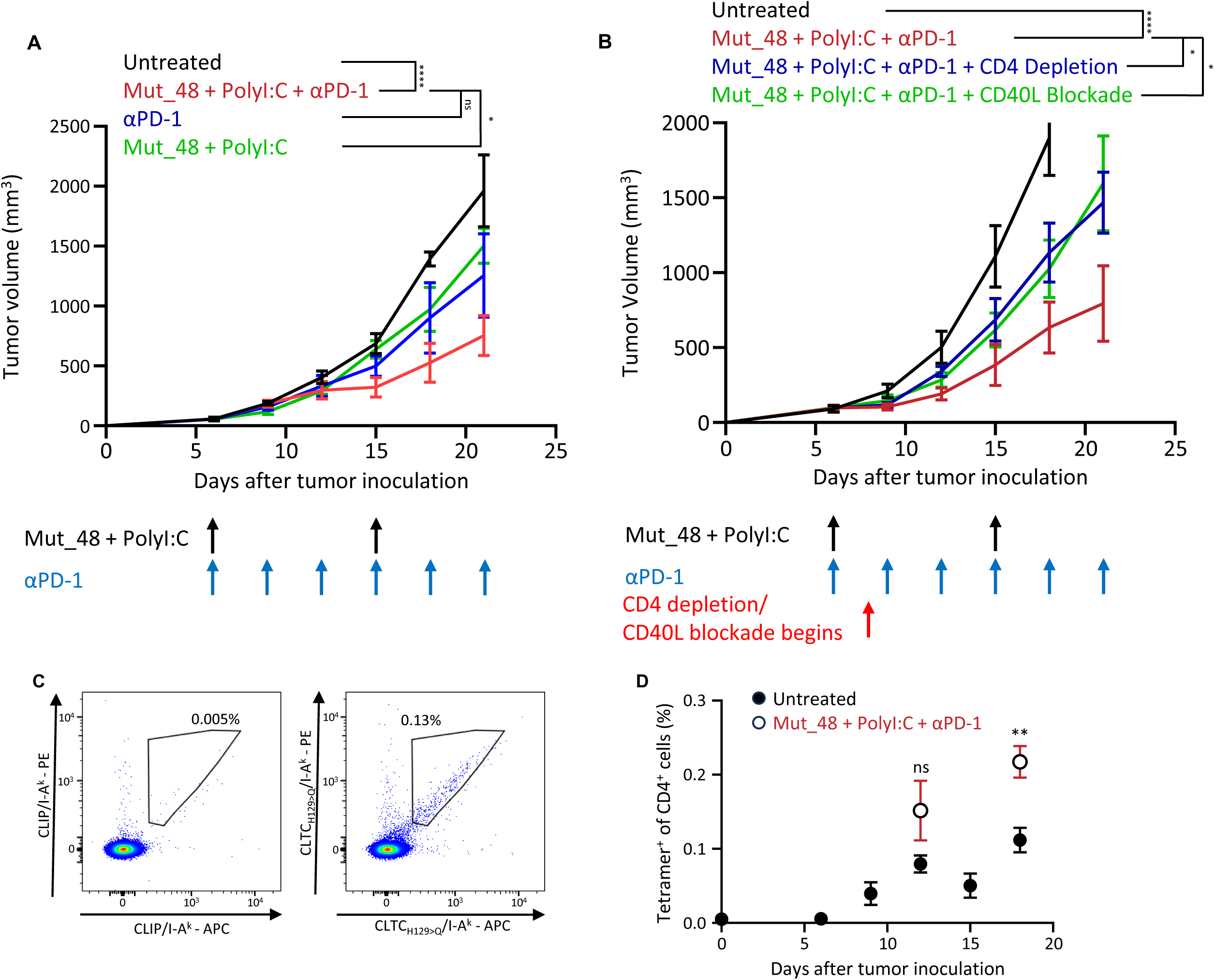
Tracking the development of NeoAg-specific CD4+ T cells against SCC VII in the tumor-draining lymph node. **a.** Tumor growth curves for mice receiving PBS (black), 100ug Mut_48 + 50ug PolyI:C + 200ug αPD-1 (red), immune checkpoint blockade alone (blue), or NeoAg vaccination alone (green) starting 6 days after SCC VII tumor challenge. n=5 for all groups. ****p < 0.0001 for Untreated vs Mut_48 + PolyI:C + αPD-1; ns p = 0.2470 for Mut_48 + PolyI:C + αPD-1 vs αPD-1; *p = 0.0434 for Mut_48 + PolyI:C + αPD-1 vs Mut_48 + PolyI:C using two-way ANOVA with Šidák correction for multiple comparisons. Data points represent mean ± s.e.m. Data depicted is from a representative experiment that was repeated with comparable results. **b.** Tumor growth curves for mice receiving PBS (black) or therapeutic vaccination plus isotype control (red), CD4 depleting (blue), or CD40/CD40L blocking (green) antibodies at the indicated time points. n=5 for all groups. ****p < 0.0001 for Untreated vs Mut_48 + PolyI:C + αPD-1; *p = 0.0258 for Mut_48 + PolyI:C + αPD-1 vs Mut_48 + PolyI:C + αPD-1 + CD4 depletion; *p = 0.0386 for Mut_48 + PolyI:C + αPD-1 vs Mut_48 + PolyI:C + αPD-1 + CD40L blockade using two-way ANOVA with Šidák correction for multiple comparisons. n=5 for all groups. Data points represent mean ± s.e.m. Data depicted is from a representative experiment that was repeated with comparable results. **c.** Representative flow cytometry plots for staining of CD4 T cells harvested from the tdLN 12 days after tumor inoculation stained with either clip (left) or CLTC /I-A^k^ tetramers (right) to identify CLTC - specific CD4^+^ T cells. **d.** Quantification of the frequency of CLTC_H129>Q_-specific CD4 T cells in the tdLN of of mice challenged with SCC VII tumors undergoing either no treatment (closed circle) or therapeutic vaccination (open circle). ns p=0.1229 for Untreated vs Mut_48 + PolyI:C + αPD-1 12 days after tumor inoculation; **p = 0.0054 for Untreated vs Mut_48 + PolyI:C + αPD-1 18 days after tumor inoculation using t-tests comparing each time point. n=4 for untreated six days after tumor inoculation and Mut_48 + PolyI:C + αPD-1 18 days after tumor inoculation. n=5 for all other points. Data points represent mean ± s.e.m. Data depicted is from a representative experiment that was repeated with comparable results.

To study the development of the CLTC_H129>Q_-specific CD4^+^ T cell response to a developing tumor, tdLN lymphocytes were isolated at 3-day intervals beginning 6 days after tumor inoculation. Staining with CLTC_H129>Q_/I-A^k^ tetramer revealed a distinct population of NeoAg-specific T cells (Figure 1c) first seen nine days after tumor inoculation which expanded throughout tumor growth without intervention, ultimately comprising approximately 0.1% of total CD4^+^ T cells in the tdLN (Figure 1d, closed symbols). CLTC_H129>Q_/I-A^k^-specific responses were contemporaneously monitored at 12- and 18-days post tumor inoculation in animals undergoing therapeutic vaccination, which approximately doubled the response’s magnitude at the later time point (Figure 1d, open symbols).

### Characterization of the CD4^+^ T cell response to SCC VII in the tdLN

Total and tetramer-positive CD4^+^ T cells were further analyzed by flow cytometry for the functional subset-defining markers CXCR3 (Th1), FoxP3 (Treg), and CXCR5^+^/PD-1^hi^ (Tfh-like). The gating strategy used to identify these cells is shown in figure 2a for total (top row) and tetramer-positive (bottom row) CD4^+^ T cells. The relative ratio of functional subsets within the total CD4^+^ T cell compartment in the tdLN showed little change over time (Figure 2b), while the CLTC_H129>Q_/I-A^k^-specific CD4^+^ T cell response was found to be comparatively diverse, involving all three functional subsets at higher frequencies than observed among total CD4^+^ T cells (Figure 2c, closed symbols). The relative frequencies of each subset varied over time and was further affected by therapeutic vaccination (Figure 2c, open symbols). NeoAg-specific Th1 cells were detected 9 days post tumor inoculation and their frequency within the tdLN rapidly declined from approximately 25% to approximately 3% of NeoAg-specific CD4^+^ cells over time regardless of therapeutic intervention. Within the same time frame, the frequency of NeoAg-specific Tregs steadily increased from 18% to 24% within the tdLN. Notably, this was reversed in animals undergoing therapeutic vaccination, which exhibited a significantly reduced frequency of NeoAg-specific Tregs of approximately 5% by day 18. CXCR5^+^ PD-1^+^ Tfh-like cells that made up a minor fraction of total CD4^+^ T cells were a significant component of the CLTC_H129>Q_-specific response, and these also subsequently deceased during progressive tumor growth.

**Fig 2:**
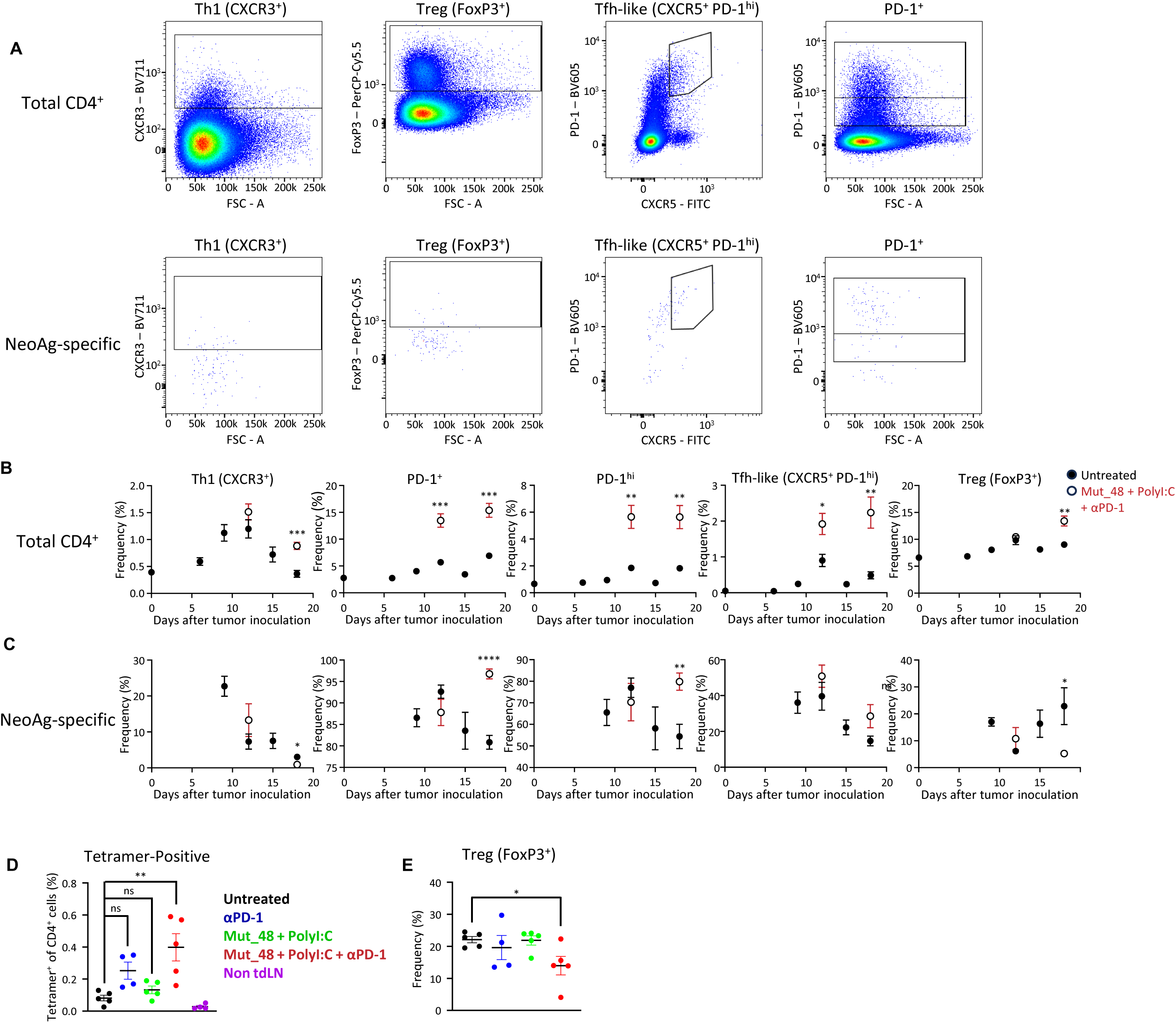
Phenotyping the developing CD4^+^ T cell response against SCC VII in the tumor-draining lymph node. **a.** Representative staining showing gating strategies for Th1, Treg, and TfH-like phenotypes as well as PD-1^+^ and PD-1^hi^ determinations on total (top row) and CLTC /I-A^k^ tetramer^+^ (bottom row) live CD4^+^ T cells isolating from the tumor-draining lymph node of tumor-bearing animals. Data obtained at 18 days after tumor inoculation. **b.** Frequencies of CXCR3, PD-1, PD-1, CXCR5 PD-1, and FoxP3 cells among total CD4 T cells in the tumor-draining lymph node throughout tumor growth without treatment (closed circles) or in mice undergoing therapeutic vaccination with Mut_48 + PolyI:C + αPD-1 (open circles). ***p = 0.0005 for CXCR3^+^; ***p=0.0004 for 12-days post tumor inoculation and ***p=0.0003 for 18-days post tumor inoculation for PD-1^+^; **p=0.0026 for 12-days post tumor inoculation and **p=0.0027 for 18-days post tumor inoculation for PD-1^hi^; *p=0.0174 for 12-days post tumor inoculation and **p=0.0044 for 18-days post tumor inoculation for CXCR5^+^ PD-1^hi^, and **p = 0.0036 for FoxP3^+^. n=5 for all groups. Data points represent mean ± s.e.m. Data depicted is from a representative experiment that was repeated with comparable results. **c.** Frequencies of CXCR3, PD-1, and PD-1, CXCR5 PD-1, FoxP3 cells among CLTC_H129>Q_-specific CD4+ T cells in the tumor-draining lymph node throughout tumor growth without treatment (closed circles) or in mice undergoing therapeutic vaccination with Mut_48 + PolyI:C + αPD-1 (open circles). *p = 0.0211 for CXCR3^+^; ****p<0.0001 for PD-1^+^; **p=0.0063 for PD-1^hi^; ns p=0.0841 for CXCR5^+^ PD-1^hi^, *p = 0.0331 for FoxP3^+^; ns p = .0572 and **p= 0.002 for PD-1^+^ of FoxP3^+^ using t-tests for each time point. n=4 for untreated 9 days after tumor inoculation. n=5 for all other points. Data points represent mean ± s.e.m. Data depicted is from a representative experiment that was repeated with comparable results. **d.** Frequencies of CLTC_H129>Q_/I-A tetramer cells among CD4 T cells taken from the tdLN 18 days after tumor inoculation in animals that received PBS (black), αPD-1 (blue), Mut_48 + PolyI:C (green), or Mut_48 + PolyI:C + αPD-1 (red) as well as from the non-draining inguinal lymph node on the opposite side of the body in animals that received Mut_48 + PolyI:C + αPD-1 (purple). ns = 0.1627 for comparing untreated to αPD-1, ns = 0.8945 for comparing untreated to Mut_48 + PolyI:C, **p = 0.0028 for comparing untreated to Mut+48 + PolyI:C + αPD-1 for using one-way ANOVA with Tukey’s correction for multiple comparisons. Data depicted is from a representative experiment that was repeated with comparable results.e. Frequency of FoxP3^+^ expression among CLTC /I-A^k^ tetramer^+^ cells observed in figure 2d. *p = 0.0298 for comparing untreated to Mut_48 + PolyI:C + αPD-1 for FoxP3^+^ frequency by t-test. Data depicted is from a representative experiment that was repeated with comparable results.

We next sought to evaluate whether the dynamics of NeoAg-specific CD4^+^ T cells and Tregs during combined immunotherapy were a result of synergistic interaction between neoantigen vaccination and immune checkpoint blockade or if either therapy arm alone was sufficient to induce a significant change. Mice were treated with α-PD-1, neoantigen vaccination, or both beginning six days after tumor inoculation. 18 days after tumor inoculation, lymphocytes were isolated from the tdLN or the non-draining inguinal lymph node on the opposite side of the body. Phenotypic analysis revealed that either therapy was insufficient to induce the significant increase in the frequency of NeoAg-specific T cells (Figure 2d) or the significant decrease in FoxP3^+^ frequency among NeoAg-specific CD4^+^ T cells (Figure 2e) in the tdLN seen with combined immunotherapy. Taken together, these results demonstrate that the CLTC_H129>Q_/I-A^k^ tetramer allows the primary NeoAg-CD4^+^ T cell response to be visualized and phenotyped within the tdLN during progressive outgrowth of SCC VII tumors. The response is both diverse and dynamic, comprising a range of functional subsets that appear with distinct kinetics and relative magnitudes. Combined immunotherapeutic intervention perturbs the NeoAg-specific response in ways that NeoAg vaccination and immune checkpoint blockade alone are unable to induce.

### Characterization of the CD4^+^ T cell response to SCC VII within the TME

To characterize the total and NeoAg-specific CD4^+^ T cells within the TME, tumor-infiltrating lymphocytes (TIL) were isolated 12- and 18-days after tumor inoculation from untreated animals and those undergoing therapeutic vaccination. In the absence of vaccine therapy, the frequency of total CD4^+^ T cells infiltrating the tumor decreased over time (Figure 3a). CLTC_H129>Q_/I-A^k^-specific cells were present within the tumor at both time points and made up a larger percentage of tumor-infiltrating CD4^+^ T cells at day 18 (Figure 3b). Most CD4^+^ T cells in the tumor had an activated phenotype, with more than 60% expressing PD-1(Figure 3c). Tregs were the dominant phenotype amongst these cells at both time points. Tregs were also the largest subset among tumor-infiltrating CLTC_H129>Q_/I-A^k^-specific cells in untreated animals, but therapeutic vaccination significantly reduced their frequency 18 days post tumor inoculation from 61% in untreated animals to 18% in vaccinated animals (Figure 3d & Supplemental Figure 1), which parallels the effect of therapeutic vaccination on CLTC_H129>Q_/I-A^k^-specific Tregs in the tdLN as seen in figure 2c. Though therapeutic vaccination did not effect the frequency of PD-1 expression on CLTC_H129>Q_-specific Tregs, it significantly reduced the frequency of PD-1 expression amongst total tumor-infiltrating Tregs from 88% to 68%.

**Fig 3:**
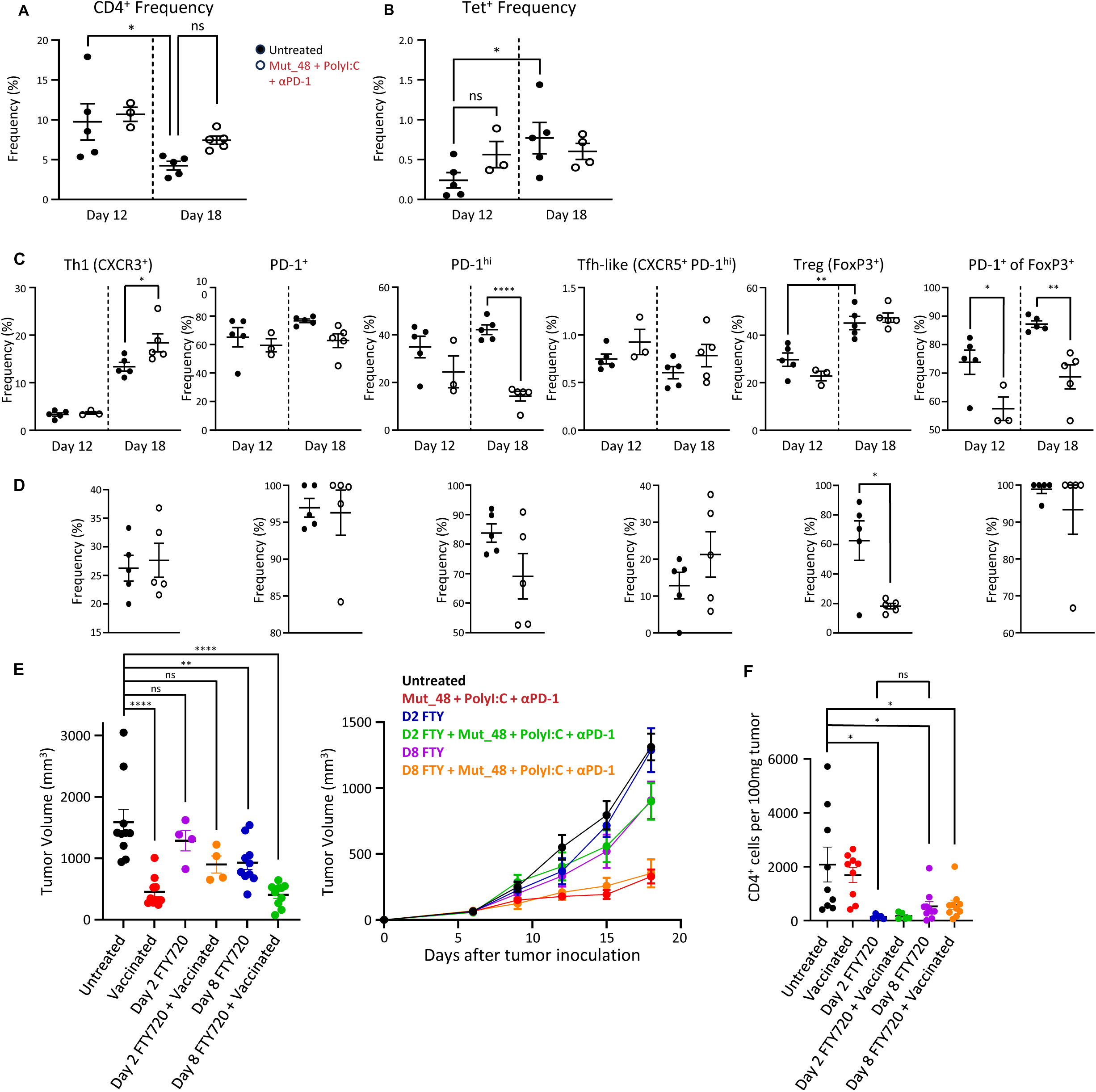
Phenotyping tumor-infiltrating CD4^+^ T cells in SCC VII. Panels a, b, c, and d compare untreated (closed circles) to animals undergoing therapeutic vaccination with Mut_48, PolyI:C, and αPD-1 as indicated in Figure 1A/Figure 1B (open circles). Data depicted is from a representative experiment that was repeated with comparable results. Bars represent mean ± s.e.m. **a.** Quantification of the frequency of CD4 T cells as a percent of all live CD45 cells isolated from the TME at the given time points. *p = 0.434; ns p = 0.3458 using one-way ANOVA with Tukey’s correction for multiple comparisons. **b.** Quantification of CLTC_H129>Q_-specific CD4 T cells as a percent of tumor-infiltrating CD4 T cells at the given time points. ns p = 0.3863; *p = 0.0458 using one-way ANOVA with Tukey’s correction for multiple comparisons. **c.** Frequencies of total CD4 cells within the tumor that are CXCR3, PD-1, PD-1, CXCR5 PD-1, FoxP3, and PD-1^+^ among Foxp3^+^ 12- and 18-days after tumor inoculation. *p = 0.0453 for CXCR3^+^; ****p<0.0001 for PD-1^hi^; **p=0.0044 for FoxP3^+^; *p= 0.0448 and **p = 0.0029 for PD-1^+^ of FoxP3^+^ using t-tests. **d.** Frequencies of CLTC_H129>Q_-specific CD4 T cells within the tumor that are CXCR3, PD-1, PD-1, CXCR5 PD-1^hi^, FoxP3^+^, and PD-1^+^ among Foxp3^+^ 18-days after tumor inoculation. *p = 0.0113 for FoxP3^+^ using t-tests. **e.** Final tumor volume for mice receiving PBS (black) or vaccinated with Mut_48, PolyI:C, and αPD-1 (red) as indicated in Figure 1a/Figure 1b with and without FTY720 administration beginning 2 (untreated in blue and vaccinated in green) and 8 days (untreated in purple and vaccinated in orange) after tumor inoculation across 2 experiments (left). ****p < 0.0001 for comparing untreated to Vaccinated, ns = 0.7908 for comparing untreated to D2 FTY, ns = 0.0542 for comparing untreated to D2 FTY + Vaccinated, **p = .0068 for comparing untreated to D8 FTY, ****p < 0.0001 for comparing untreated to D8 FTY + Vaccinated using one-way ANOVA with Tukey’s correction for multiple comparisons. Tumor growth curve from 1 experiment (right) that was repeated with comparable results. Bars represent mean ± s.e.m. **f.** Quantification of CD4 T cells found per 100mg of tumor in untreated mice (black) and animals undergoing immunotherapeutic vaccination using Mut_48, Poly(I:C), and αPD-1 beginning 6 days after tumor inoculation (red circles) with and without FTY720 administration beginning 2- and 8-days after tumor inoculation. *p = 0.0129 for comparing untreated to D2 FTY720, *p = 0.0172 for comparing untreated to D8 FTY720, *p = 0.0237 for comparing untreated to D8 FTY720 + vaccinated, ns = 0.9784 for comparing D2 FTY720 to D8 FTY20. Data shown is from 2 experiments.

We next investigated the temporal dynamics of lymphocyte egress from secondary lymph tissue by administering FTY720 daily beginning either 2- or 8-days post tumor inoculation in untreated animals and in those receiving Mut_48, PolyI:C, and αPD-1. FTY720 treatment beginning 2 days but not 8 days post tumor inoculation prevented immunotherapeutic vaccination from significantly reducing tumor burden (Figure 3e) even though any group that received FTY720 had significantly fewer CD4^+^ T cells within the tumor compared to animals that did not (Figure 3f).

These data reveal key similarities and differences between the NeoAg-specific CD4^+^ T cell response within the tumor and draining lymph node throughout the course of tumor development, with the early appearance and increasing presence of *Cltc*-specific Tregs in both locations offering possible clues into how an initial Th1 response is followed by a potent population of NeoAg-specific regulatory cells.

### Single-cell genomic analysis of the natural CLTC_H129>Q_-specific CD4^+^ T cell response

We sought to better understand key features of the spontaneous CLTC_H129>Q_/I-A^k^-specific response that may explain its dynamic nature by comparing the transcriptional profiles and TCR repertoire of single cells. Figure 4a diagrams the approach used to obtain single-cell RNA and TCR data through the 10X platform. Global gene expression profiling of sorted CLTC_H129>Q_/I-A^k^ tetramer-positive versus tetramer-negative cells showed that these populations are remarkably distinct, with the majority of tetramer-negative cells expressing a naïve phenotype (Figure 4b, Supplemental Figure 2a, and Supplemental Figure 2b). Tetramer-positive cells, in contrast, were antigen-experienced and had differentiated into specific functional subsets (Figure 4c) including stem-like cells with high levels of TCF1 and few markers of differentiation, T_H_1 cells that express higher levels of Ifng and TNF, regulatory cells with high levels of IL-10, Lag-3, and IL-2ra, and T_FH_-like cells expressing high levels of IL-6, CXCR5, and Bcl6 (Figure 4d). The relative frequencies of the regulatory and Tfh-like clusters observed is similar to that identified by flow cytometry (Supplemental Figure 2c), though the frequency of the Th1 cluster was notably higher than when observed by flow cytometry. Vβ family analysis indicated a higher degree of clonal expansion in tetramer-positive than in tetramer-negative cells (Figure 4e). These data demonstrate a remarkably high degree of phenotypic heterogeneity present at the single cell level within the endogenous primary NeoAg-specific CD4^+^ T cell response to a developing tumor.

**Fig 4:**
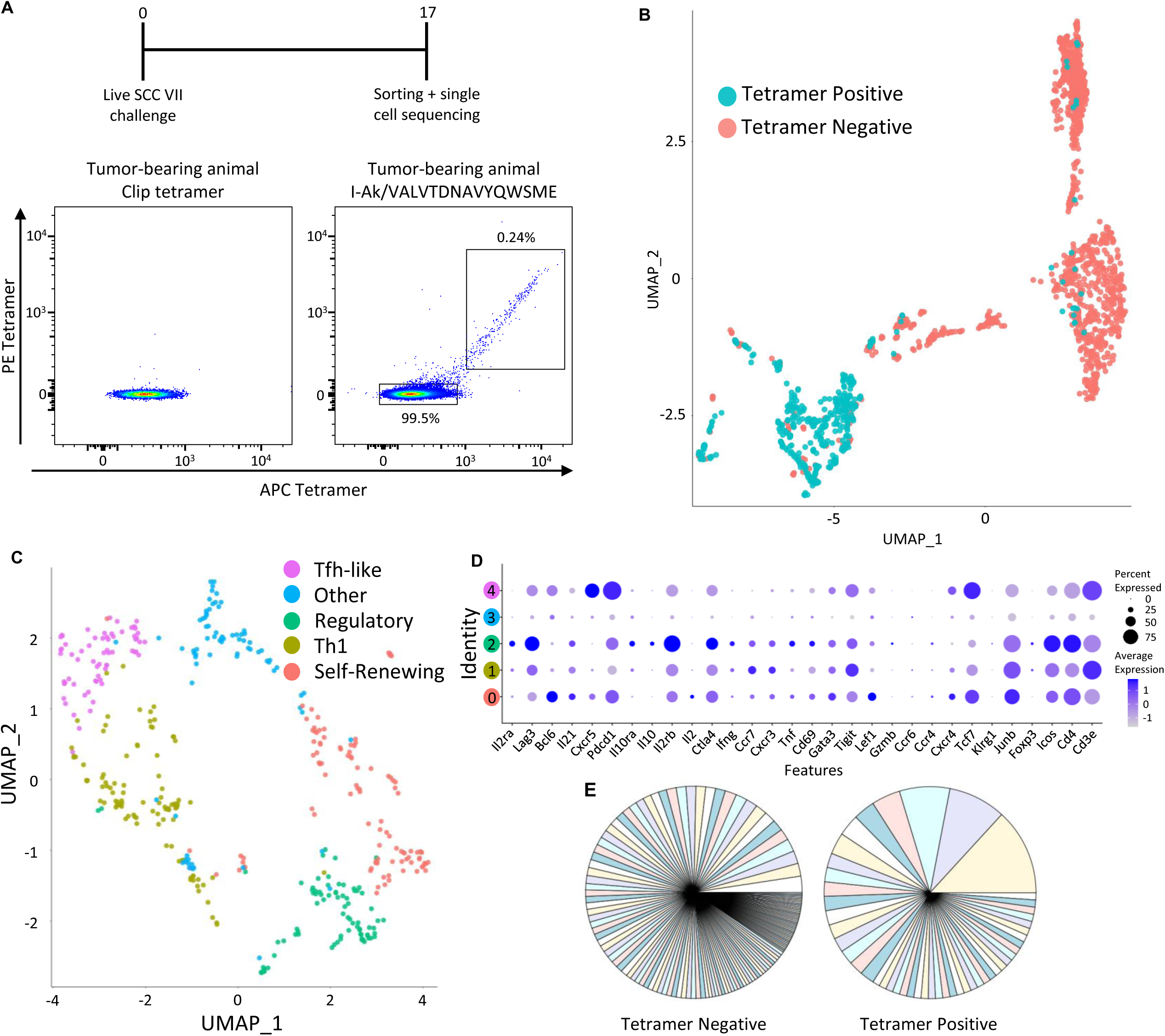
Investigating the clonality and genetic profile of the neoantigen-specific CD4^+^ T cell response using single cell genomics. **a.** Schematic overview and flow cytometry plots of tetramer sorting experiment to isolate CLTC_H129>Q_-specific CD4^+^ T cells from tumor-draining lymph nodes (top) plus tetramer staining used to sort tetramer-negative and tetramer-positive cells for further analysis. **b.** The UMAP of CLTC_H129>Q_/I-A tetramer positive (blue) and tetramer negative (pink) cells determined by Seurat-based clustering. There were a total of 1,581 cells, 432 of which are tetramer-positive and 1,149 of which are tetramer-negative. **c.** The UMAP of tetramer positive cells as determined by Seurat-based clustering. A total of 5 clusters (clusters 0-4) were identified and color-coded. **d.** The average expression of subset-defining and functional genes by cluster pictured in panel 5c. **e.** TCR Vβ gene diversity from sorted CLTC_H129>Q_/I-A tetramer negative and tetramer positive cells.

### Developmental plasticity within the Cltc_H129>Q_-specific TCR repertoire

We next examined the phenotype of CD4^+^ T cells expressing distinct NeoAg-specific TCRs. Individual α/β TCR clonotypes isolated from CLTC_H129>Q_/I-A^k^-binding cells (Table 1) were expressed in primary C3H CD4^+^ T cells by retroviral transduction using a variant of the pMSGV1 vector diagramed in figure 5a encoding TCR genes upstream of CD90.1, which was used as a transduction reporter. Clonotypes that successfully bound CLTC_H129>Q_/I-A^k^ tetramer (Table 2) were then mapped back to the differential gene analysis for quantification of their clustering (Figures 5b and 5c). This analysis revealed that nearly every TCR was found in multiple clusters, with no clear preference for any individual TCR to limit the range of fates it was able to adopt. These findings demonstrate that, rather than being constrained to a specific functional subset, different developmental fates are available to cells expressing the same clonal TCR.

**Fig 5:**
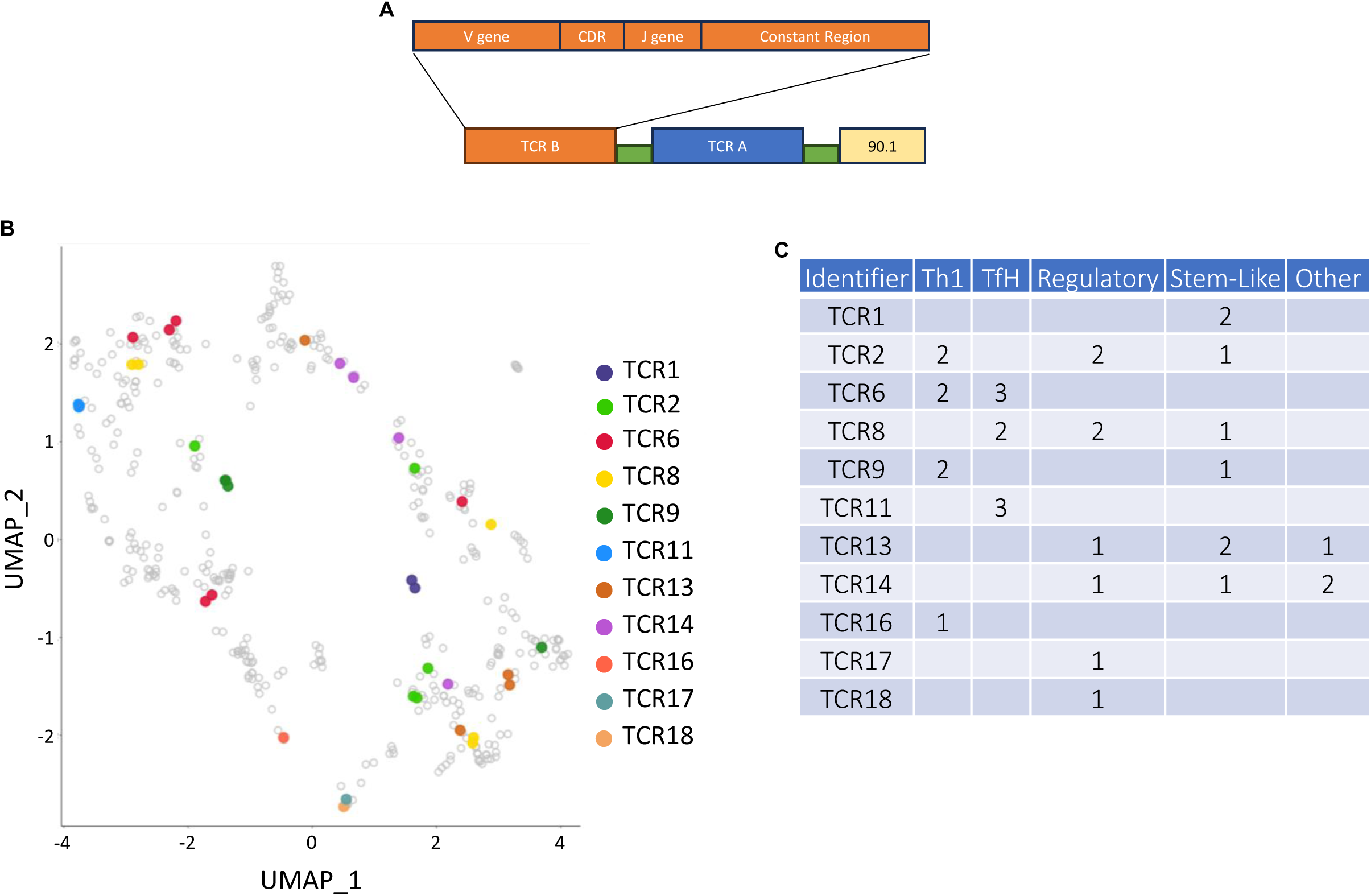
Characterizing the clonality and phenotypes of CLTC_H129>Q_-specific TCRs. **a.** Schematic construct for retroviral production and transfection containing TCR genes of interest alongside CD90.1. **b.** Confirmed NeoAg-specific TCRs overlayed on the UMAP seen in figure 4c. **c.** Quantification of the number of cells for each CLTC_H129>Q_-specific clone found within the given clusters determined in figure 4c and figure 4d.

### Functional characterization of the Cltc_H129>Q_--specific TCR repertoire

We next investigated whether the affinity of a given TCR influences the developmental fate of the T cells expressing it. This was done by titrating the concentration of tetramer used during staining (Figure 6a). CD90.1 was used as a surrogate for TCR expression as Vβ staining was also observed on some percent of non-transduced cells naturally expressing the given Vβ gene (Supplemental Figure 3). We normalized differences in retroviral transduction efficiency between TCRs using a ratio of the mean fluorescent intensity of CLTC_H129>Q_-tetramer and the CD90.1 reporter (Figure 6b). The 11 distinct validated NeoAg-specific TCRs have a wide range of affinities, including a 17.75 fold range at the standard working tetramer concentration of 2µg/mL used in previous experiments (Figure 6c). We then mapped the affinity of each TCR back to the UMAP used to determine phenotypic clustering in Figure 4c (Figure 6d). This revealed that each functional subset was adopted by TCRs with a wide range of affinities without any correlation between TCR affinity and the phenotypes adopted. (Figure 6e).

**Fig 6:**
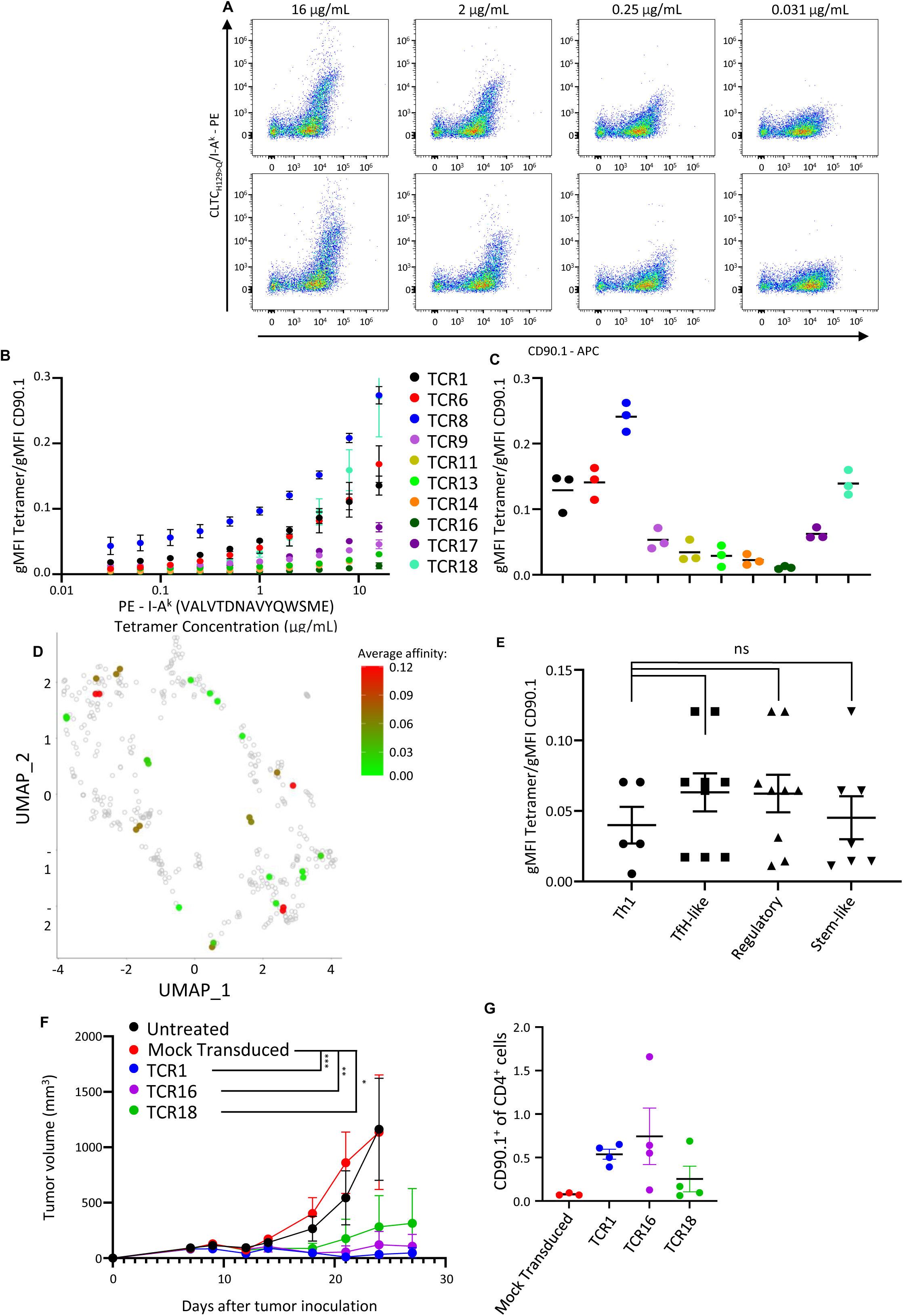
Expression and characterization of NeoAg-specific TCRs. **a.** Flow cytometry plots showcasing the titration of CLTC_H129>Q_/I-A tetramer on two different TCRs with similar affinities. **b.** Quantification of tetramer affinity of validated neoantigen-specific TCRs normalized using CD90.1 over the range of concentrations tested during titration. All data is from 3 experiments. Points represent mean ± s.e.m. **c.** Quantification of normalized tetramer affinity for validated neoantigen-specific TCRs at a concentration of 2ug/mL used throughout figures 1, 2, and 3. All data is from 3 experiments. Points represent mean ± s.e.m. **d.** Heat map of normalized mean tetramer affinity determined in figure 6c overlaid on the UMAP seen in figure 4c. **e.** Quantification of TCR affinity for clones within each of the phenotypic clusters determined in figure 4c. p = 0.7073 for Th1 vs TfH-like; p = 0.7299 for Th1 vs Regulatory; p = 0.9955 for Th1 vs Stem-like using one-way Anova. Bars represent mean ± s.e.m. **f.** Tumor growth curves for mice that received cyclophosphamide seven days post tumor inoculation and either PBS (black) or 3 x 10^6^ adoptively transferred CD4^+^ T cells that underwent mock transduction (red) or were transduced to express TCR1 (blue), TCR16 (purple), or TCR18 (green) eight days after tumor inoculation. ***p = 0.0007 for Mock Transduced vs TCR1; **p = 0.0033 for Mock Transduced vs TCR16; *p = 0.0352 for Mock Transduced vs TCR18 using two-way ANOVA with Šidák correction for multiple comparisons. n=4 for all points. Data points represent mean ± s.e.m. Data depicted is from a representative experiment that was repeated with comparable results. **g.** Frequency of CD90.1 cells as a percent of all CD4 T cells found within the tumor-draining lymph node in animals treated with cyclophosphamide and adoptive transfer of CD4^+^ T cells that were either mock transduced (red) or were transduced with TCR1 (blue), TCR16 (purple), or TCR18 (green) as described in panel 6f. Cells were harvested 27 days after tumor inoculation. Bars represent mean ± s.e.m. Data depicted is from a representative experiment that was repeated with comparable results.

To further demonstrate that TCR affinity does not determine phenotypic fate or function, we asked whether TCRs isolated from Tregs could nonetheless mediate therapeutic efficacy when used for ACT. To examine this, primary CD4^+^ T cells were transduced with TCR1, a high-affinity TCR which we have previously shown can mediate tumor rejection in the context of ACT, or with either TCR16 or TCR18, each of which were isolated from a single FoxP3^+^ cell. Following TCR engineering, the transduced cells were cultured with IL-7 and IL-15 to induce a Tscm phenotype prior to adoptive transfer into cyclophosphamide-conditioned C3H mice bearing established SCC VII tumors inoculated 8 days prior. Despite a 12.67-fold range of affinities between the distinct TCRs and a 3.95-fold difference in transduction efficiency (Supplemental Figure 4), all three TCRs were able to successfully clear tumors in some animals and reduce tumor burden in the rest while transfer of mock transduced polyclonal cells had no discernable effect (Figure 6f). Additionally, CD90.1^+^ cells were found at comparable levels in the tdLN 19 days after injection without any apparent correlation between the TCR they expressed and either the frequency of CD90.1^+^ cells or the residual tumor burden (Figure 6g).

## Discussion

Tumor-specific CD4^+^ T cell responses have traditionally been studied in the ‘post-hoc’ setting following their generation by targeted vaccination, immune checkpoint blockade, or at late stages of tumor outgrowth. Although such studies have been instrumental in revealing the various effector and regulatory roles of CD4^+^ T cells in tumor immunity, none have allowed insights into the natural ontogeny of tumor-specific CD4^+^ T cells in unmanipulated conditions. The availability of a CLTC_H129>Q_/I-A^k^ tetramer has enabled the characterization of the natural CD4^+^ T cell response to a validated NeoAg for the first time. We investigated progressively growing tumors and those treated with therapeutic vaccination to study the TCR repertoire, phenotypic fates, and relative functional avidity of different subsets of NeoAg-specific CD4^+^ T cells.

The results of these analyses reveal that the endogenous CD4^+^ T cell response to a cross-presented MHC-II-restricted neoantigen is both diverse, involving differentiation into several distinct functional subsets, and dynamic, with the relative frequencies of regulatory and different effector subsets changing over time with progressive tumor growth and in response to therapeutic vaccination.

Our results show that the CD4^+^ T cell subsets comprising the CLTC_H129>Q_/IA-k-specific response is dynamic. This study used flow cytometry for initial phenotyping, single-cell transcriptomics to verify the gene expression programs that characterize the diverse CLTC_H129>Q_/I-Ak-specific CD4^+^ T cell response, and applied TCR engineering to probe the TCR repertoire. These results echoed the diversity found in our multimodal phenotypic analysis and show that the initial response within the tdLN is comprised of Th1, Tfh-like, and Treg cells expressing TCRs spanning a range of receptor affinities (Figure 2c and 4d).

The frequency of NeoAg-specific Th1 cells decreases in the tdLN throughout tumor growth. Therapeutic vaccination with the mutated CLTC peptide, polyI:C, and α-PD-1 significantly reduces both the frequency and total number of NeoAg-specific Tregs in both the tdLN and TME 18 days after tumor inoculation (Figure 2c and 3d). Given the early rise of NeoAg-specific Tregs and that therapeutic vaccination did not reduce the frequency of total Tregs, the reduction in NeoAg-specific Tregs at later time points seems to be specific to the antigen included in the vaccine. It appears that vaccination directly influences early-migrating CD4^+^ T cells in the TME without requiring cells from the tdLN, as using FTY720 to block exfiltration of lymphocytes from the tdLN beginning 2- but not 8-days after tumor inoculation reduced the efficacy of immunotherapeutic intervention (Figure 3e).

Our data clearly show that the CLTC_H129>Q_/I-A^k^-specific CD4^+^ T cell response is oligoclonal and comprised of TCRs exhibiting a range of affinities. However, TCR affinity did not appear to correlate with the phenotypic fates of each clone (Figure 6e). We adoptively transferred primary CD4^+^ T cells retrovirally-transduced to express a NeoAg-specific TCR found during the endogenous response. All tested NeoAg-specific TCRs, including the lowest affinity TCR identified as being expressed solely by a single FoxP3^+^ cell, were capable of mediating tumor clearance to a comparable extent and were found within the tumor-draining lymph nodes at similar frequencies independent of the functional subset from which the TCR was identified, TCR affinity, or transduction efficiency. Given that Tregs can be found amongst NeoAg-specific cells at the earliest time points the response can be identified, this raises the interesting possibility that a therapy capable of converting these early-rising NeoAg-specific Tregs into anti-tumor phenotypes would be extremely potent.

There is mixed evidence suggesting that the affinity of CD8^+^ T cell TCRs for their cognate antigen influences their fate or therapeutic potential (28, 29). The critical role for cytokines in determining the fate of CD4^+^ T cells lends credence to the idea that TCR affinity plays a limited or negligible role. Given the diversity of the initial NeoAg-specific CD4^+^ T cell repertoire as soon as the response is able to be detected, it’s unclear if the changes to the relative ratios of phenotypic subsets throughout tumor growth are a result of changing conditions for initial priming of newly activated clones or a result of the environments, APC types available, and accessory signals inducing either preferential division in a limited subset of phenotypes or plasticity in the ongoing response.

These findings offer the first look into the form and functionality of a developing natural NeoAg-specific CD4^+^ T cell response. Our data reveals the range of functional subsets that the response can comprise and demonstrates how combined neoantigen therapy and αPD-1 immune checkpoint blockade may alter these and reduce tumor burden. Additionally, our study shows the diversity of the response to a single neoantigen can generate clonal T cells which can adopt a range of developmental fates that are independent of their TCR avidity. These insights will aid in understanding the diversity and functionality of antitumor CD4^+^ T cell responses and should allow the development of strategies through which these can be therapeutically manipulated to favor tumor control.

## Methods

### Animals

Male and female C3H/HeJ mice (The Jackson Laboratory, Bay Harbor, ME) were used in these experiments. Animals used were 7-12 weeks of age and maintained/bred in the La Jolla Institute for Immunology vivarium with pathogen-free conditions in accordance with guidelines of the Association for Assessment and Accreditation of the Laboratory Animal Care International.

### Cell Culture

Squamous cell carcinoma VII San Francisco (SCC VII) is a tumor line which spontaneously arose in the abdominal wall of a C3H mouse at the laboratory of Herman Suit (Harvard University, Cambridge, MA) and was adapted for *in vitro* growth by Karen K. Fu and Kitty N. Lam (University of California, San Francisco, CA). SCC VII was maintained for a maximum of 3 passages *in vitro* in RMPI 1640 growth medium with 12.5% fetal bovine serum (FBS) and 100 units/mL each of penicillin/streptomycin (Gibco). Vials of P0 SCC VII cells were obtained by injecting female C3H/HeJ mice subcutaneously (S.C.) with 5×10^5^ P2 cells in 1x DPBS and harvesting tumors 14 days after inoculation. Tumor tissue was dissociated with the mouse Tumor Dissociation Kit (Miltenyi Biotec, Aubrun, CA), passed through a 70µm cell strainer (Fisher Scientific, Pittsburgh, PA), and resuspended in media for *in vitro* passage.

### Injections & in vivo treatments

SCC VII tumors were generated by injecting 5×10^5^ SCC VII cells in 100µL 1x DPBS S.C in the left flank. Peptide vaccinations were carried out 6 and 15 days post tumor inoculation in the parental SCC VII line by injecting 100µg peptide and 50µg Poly(I:C) combined in 100µL of 1x DPBS S.C. directly anterior to the growing tumor. Immune checkpoint blockade was carried out by intraperitoneal (I.P.) injection of 200µg of αPD-1 (Clone RMP1-14, Bio X Cell) compared to polyclonal rat IgG2b κ (Bio X cell) as an isotype control every 3 days starting concurrently with peptide vaccination. CD4 depletion was achieved by I.P. injection of 400ug αCD4 (Clone GK1.5, in house) compared to polyclonal rat IgG2b κ (Bio X cell) as an isotype control 9 days after tumor inoculation. CD40/CD40L was blocked by I.P. injection of 200ug αCD40L (MR1, Bio X Cell) compared to polyclonal Aremnian hamster IgG (Bio X Cell) isotype control every 2 days beginning 9 days after tumor inoculation.

Where indicated, tumor-bearing C3H/HeJ mice were treated with 3mg (150mg kg^-1^) cyclophosphamide monohydrate (Sigma) dissolved in 1x PBS i.p. 7 days post tumor inoculation. Tumor-bearing Cy-treated animals were then injected i.v. via the tail vein with 3 million CD4^+^ T cells in 200 uL HBSS 8 days post tumor inoculation.

Tumor volume was calculated using the equation V = L*W^2^/2 where L represents the longest length of the tumor and W represents the width perpendicular to the longest length. Prior to the first round of treatments, tumors were measured and groups were normalized to equalize starting tumor volume with outliers removed to reduce variability. N=5 was chosen for animal experiments to balance cost and power. Cages were treated and measured in a random order to minimize potential confounders. Experiments were not blinded as one person handled all treatments and measurements.

### Preparation of single cell suspensions

Lymph nodes were surgically removed and dissociated manually via passage through a 70µm strainer. Tumors were surgically removed and cut into small (<2mm) pieces. The fragments were enzymatically dissociated in 20 µg/mL Liberase (Roche) and 20 µg/mL DNase I (Roche) at 37 degrees for 45 minutes. They were shaken at the 15 and 30 minute marks. Afterwards, they were passed through a 70 µm strainer.

### Flow cytometry

Prior to staining, Fc receptors were blocked with αCD16/32 (Clone 2.4G2, BD Biosciences) for 15 minutes at 4C. Tetramer staining was carried out using PE- or APC-conjugated I-A^k^(VALVTDNAVYQWSME) or CLIP tetramer PVSKMRMATPLLMQA. They were incubated at the indicated concentration in FACS buffer (PBS + 1% bovine serum albumin + 0.1% sodium azide) at 37C for 1 hour. Surface antigens were stained with the indicated antibodies for 20 minutes on ice. Dead cells were excluded using either DAPI immediately prior to reading or Fixable Viability Dye eFlour 780 (ThermoFisher) prior to surface staining. Transcription factor staining was carried out using the eBioscience Foxp3 / Transcription Factor Staining Buffer Set (eBioscience) according to manufacturer guidelines. All flow cytometry was performed on a BD FACS Celesta or BD LSR-II and analyzed using FlowJo.

### Single cell genomics

Single-cell (sc)RNA-seq reads were demultiplexed into individual libraries (n=12). Demultiplexing was executed using Illumina’s bcl2fast software (v2.19.0.316). The individual libraries underwent processing with 10x Genomic’s Cell Ranger software (3.1.0). The module count was employed to align and map the reads to the mm10 reference mouse genome (v3.0.0). Unique molecular identifier (UMI) counts for each cell barcode associated with each gene. Gene expression data from the individual libraries were aggregated into a single dataset that included both tetramer-positive and tetramer-negative cells.

The Seurat toolkit (v3.1.5) was employed to conduct the analysis of the aggregated scRNA-seq dataset. Genes expressed by less than 0.1% of the cells in the dataset were excluded from the analysis. T-cell receptor TCR (annotations) were performed as a distinct step (described below). To reduce the occurrence of potential doublets and transcriptomes of low quality, the barcodes that did not meet the following criteria were filtered out: UMI counts between 500 and 15,000, gene counts between 300 and 4,500, and a mitochondrial UMI percentage not exceeding 10%.

The data of the aggregated dataset was normalized using the NormalizeData function in Seurat (default parameters). The most variable genes were identified as those with a mean UMI capture greater than 0.01, contributing to 25% and 10% of the total standardized variance in the aggregate dataset. This selection process was carried out by applying the variance-stabilizing transformation method through the function FindVriableFeatures (default parameters). Data scaling was performed using the ScaleData function, with custom variables regressed out using the ‘vars.to.regress’ parameter. Cell clusters were determined within the aggregated dataset using the FindNeighbors and FindClusters functions, primarily with default parameters, except for the number of principal components (PCs; PCs=20) and the ‘resolution’ parameter (res). The RunUMAP function was applied with the top 40 PCs, 15 neighboring points for local approximations of the manifold structure, and a minimum distance of 0.1 (specified as arguments dims, n.neighbors, and min.dist, respectively) to compute manifold approximation and projection (UMAP) embeddings. Various plot types were generated using custom scripts, primarily based on ggplot2, an open-source data visualization package for the R statistical programming language, and are available from the GitHub repository (Github repo).

### TCR cloning and expression

TCR amino acid sequences were constructed from single cell genomics data. DNA sequences were generated using NovoPro ExpOptimzer. The sequences were synthesized and cloned into MSGV1 retroviral expression backbones using BioXP (Codex DNA, San Diego CA). TCR alpha and beta chains were separated from each other and a flanking CD90.1 sequence with a P2A ribosomal skipping element. TCR retroviral supernatants were harvested 48 and 72 hours after cotransfection of Platinum-Eco cells with the TCR-containing retroviral vectors and pcL-Eco. Supernatants were stored at −80C. For transfection of mouse primary cells, spleens from naïve C3H/HeJ mice were manually dissociated. Cell suspensions were passed through a 70 µm filter. CD4^+^ T cells were isolated by magnetic negative selection (StemCell) then stimulated for 24 hours with anti-CD3/CD28 Dynabeads (Gibco) in T cell media (RPMI 1640 + 10% FBS + 50µM beta-mercaptoethanol, 1x, penicillin/streptomycin and HEPES). Non-tissue Culture-Treated 24 well plates (Falcon). were coated with 10µg/mL retronectin overnight at 4C. They were washed with 2% BSA in HBSS for 30 minutes at room temperature, then retroviral supernatants were spun down at 2000g for 2 hours at 32C. After discarding the supernatant, 1 million activated mouse primary cells were spun down in each well at 500g for 10 minutes with break and acceleration of 1.

## Declarations

All animal work undertaken in this study was performed under protocol number AP007-SPS1 as approved Consent for publication.

All authors listed have given their consent for publication

All data, T cell receptors, and cell lines used or generated in this study will be made available to academic colleagues upon request,

This work was funded by NIH UO1 DE028227 to SPS

The contributions of each author are as follows: Concept and experimental plans: RG. SPS, AM, EC experimental procedures: RG, KSZ, ND, MN, RT, HD, SA, Data analysis: RG, SPS, MAO, PV, BP. Manuscript writing: RG, SB, SPS.

## Acknowledgements

The authors wish to thank the leadership and staff of the LJI Department of Laboratory Animal Care, Flow Cytometry Core Facility, and Next-Generation Sequencing Core Facility without whom this work could not have been performed, and tp the National Institutes of Health and National Cancer Institute for the funding support.

---

**Supplemental figure 1: Therapy decreases the frequency of NeoAg-specific FoxP3^+^ cells**

**a.** Representative FACS plots of tumor-resident tetramer-positive CD4+ T cells from untreated and vaccinated animals taken 18 days after tumor inoculation.

**Supplemental Figure 2: Different activation states for total and CLTC_H129>Q_-specific cells.**

**a.** UMAP with grouping based on activation states determined by Seurat-based clustering originally pictured in figure 4b. Group 0 is primarily composed of tetramer positive cells while groups 1 and 2 are composed of tetramer negative cells.

**b.** Relative CD44 expression of the three groups seen in panel a.

**c.** The frequency of each subset identified 18 days after tumor inoculation through flow cytometry in panel 2c and 17 days after tumor inoculation through single cell genomics in figure 4c

**Supplemental Figure 3: CD90.1 vs TCR expression**

**a.** Flow plots comparing CD90.1 to the TCRVß gene contained in the TCR1 construct in CD4+ T cells that underwent either mock transduction (left) or that were transduced with the construct containing TCR1 (right).

**Supplemental Figure 4: Transduction efficiency of adoptively transferred cells.**

**a.** CD90.1 expression versus tetramer binding for the four sets of adoptively transferred cells used in figures 6f and 6g analyzed before injection

**Table 1: List of the TCRs**

**Table 2: Expression levels and tetramer binding by TCRs**

