## Supplementary Figures for "The spontaneous neoantigen-specific CD4^+^ T cell response to a growing tumor is functionally and phenotypically diverse"

### Supplementary Figure 1

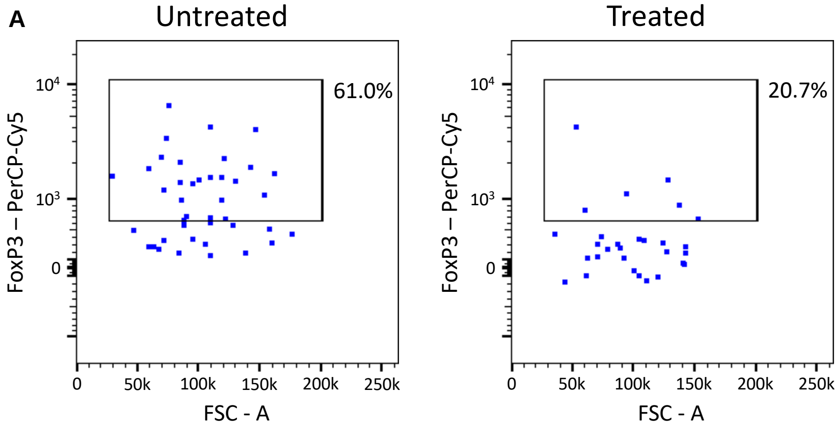

Supplementary Figure 2

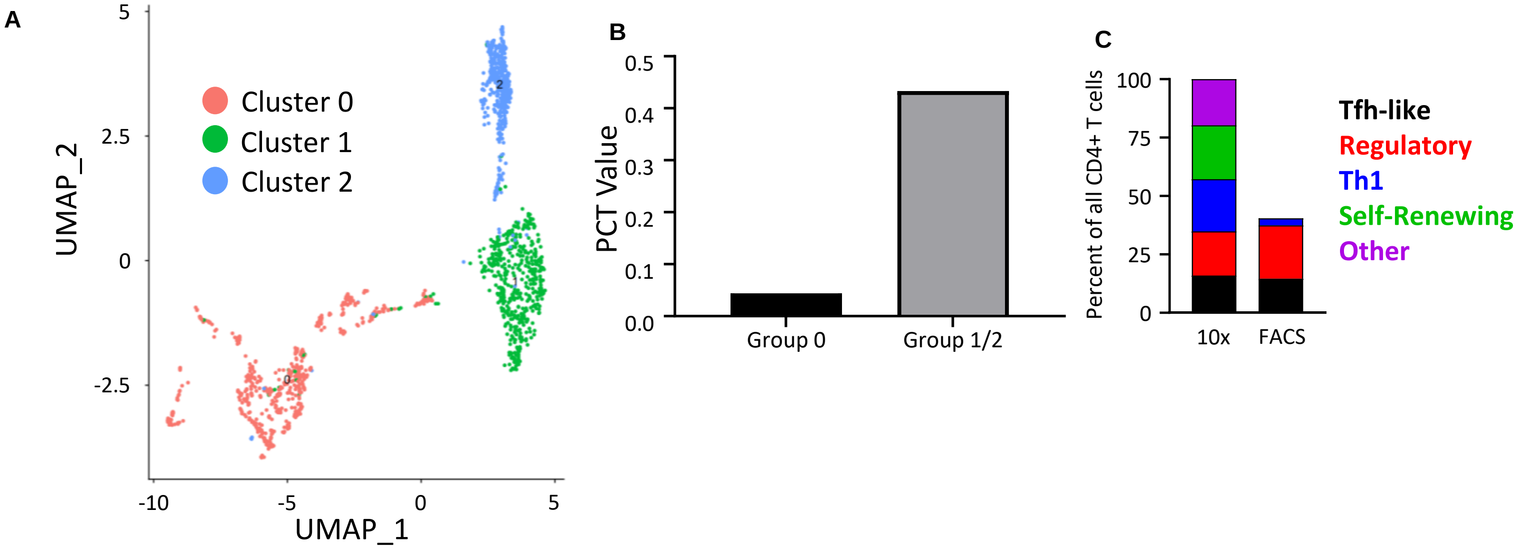

### Supplementary Figure 3

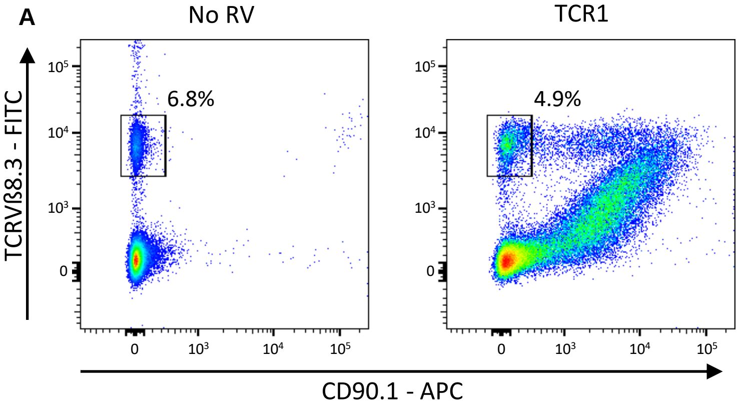

Supplementary Figure 4

**A**

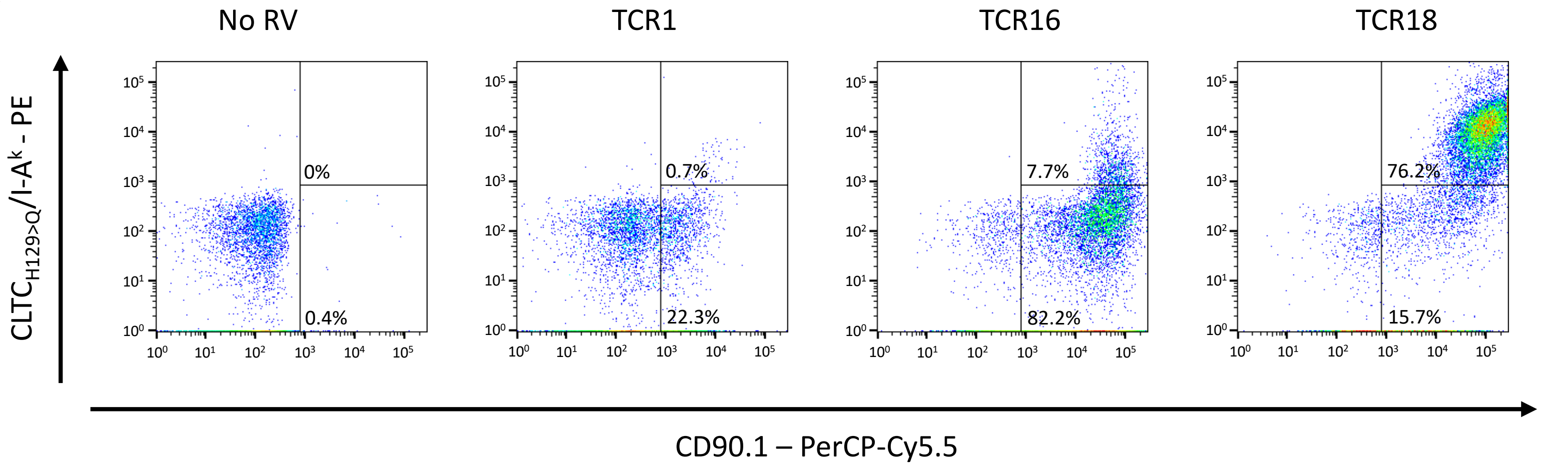

Table 1

| Clone ID | Clone # | TRAV | TRAJ | TRBV | TRBJ |
|---|---|---|---|---|---|
| TCR1 | 2 | TRAV9-4 | TRAJ5 | TRBV13-1 | TRBJ1-1 |
| TCR2 | 5 | TRAV9-4 | TRAJ5 | TRBV13-1 | TRBJ1-1 |
| TCR3 | 11 | TRAV13-4-DV7 | TRAJ27 | TRBV10 | TRBJ1-4 |
| TCR4 | 9 | TRAV14D-1 | TRAJ21 | TRBV13-3 | TRBJ1-2 |
| TCR5 | 8 | TRAV4-2 | TRAJ26 | TRBV1 | TRBJ2-5 |
| TCR6 | 6 | TRAV7D-5 | TRAJ15 | TRBV2 | TRBJ1-5 |
| TCR7 | 6 | TRAV7D-4 | TRAJ7 | TRBV5 | TRBJ2-5 |
| TCR8 | 5 | TRAV7-1 | TRAJ37 | TRBV2 | TRBJ2-5 |
| TCR9 | 5 | TRAV6D-4 | TRAJ43 | TRBV12-2 + TRBV13-2 | TRBJ1-4 |
| TCR10 | 5 | TRAV3-4 | TRAJ40 | TRBV12-2 + TRBV13-2 | TRBJ2-7 |
| TCR11 | 4 | TRAV7D-5 | TRAJ37 | TRBV2 | TRBJ2-1 |
| TCR12 | 4 | TRAV14D-1 | TRAJ21 | TRBV13-3 | TRBJ1-4 |
| TCR13 | 4 | TRAV14D-2 | TRAJ9 | TRBV31 | TRBJ1-3 |
| TCR14 | 4 | TRAV4-3 | TRAI9 | TRBV1 | TRABJ1-3 |
| TCR15 | 5 | TRAV14D-2 | TRAJ40 | TRBV1 | TRBJ2-7 |
| TCR16 | 1 | TRAV6N-7 | TRAJ57 | TRBV2 | TRBJ2-7 |
| TCR17 | 1 | TRAV7-1 | TRAJ37 | TRBV2 | TRBJ2-7 |
| TCR18 | 1 | TRAV7-3 | TRAJ15 | TRBV2 | TRBJ2-7 |
| TCR19 | 7 | TRAV4-2 | TRAJ37 | TRBV20 | TRBJ1-6 |
| TCR20 | 6 | TRAV6-2 | TRAJ6 | TRBV20 | TRBJ2-7 |

Table 2

| Clone ID | Clone # | CD90.1 Expression | Tetramer Binding |
|---|---|---|---|
| TCR1 | 2 | Yes | Yes |
| TCR2 | 5 | Yes | Yes |
| TCR3 | 11 | Yes | No |
| TCR4 | 9 | Yes | No |
| TCR5 | 8 | Yes | No |
| TCR6 | 6 | Yes | Yes |
| TCR7 | 6 | Yes | No |
| TCR8 | 5 | Yes | Yes |
| TCR9 | 5 | Yes | Yes |
| TCR10 | 5 | Yes | No |
| TCR11 | 4 | Yes | Yes |
| TCR12 | 4 | No | No |
| TCR13 | 4 | Yes | Yes |
| TCR14 | 4 | Yes | Yes |
| TCR15 | 5 | Yes | No |
| TCR16 | 1 | Yes | Yes |
| TCR17 | 1 | Yes | Yes |
| TCR18 | 1 | Yes | Yes |
| TCR19 | 7 | Yes | No |
| TCR20 | 6 | Yes | No |
